# Genetic interaction of *Specc1l* and *Thm1* reveals cytoskeletal–ciliary crosstalk

**DOI:** 10.1101/2025.11.03.686369

**Authors:** Brittany M. Hufft-Martinez, Dana N. Thalman, An J. Tran, Jeremy P. Goering, Marta Stetsiv, Michael Moedritzer, Sarah C. Wilson, Henry H. Wang, Pamela V. Tran, Irfan Saadi

**Author notes:** Correspondence should be addressed to: Irfan Saadi, PhD, Department of Cell Biology and Physiology, University of Kansas Medical Center, 3901 Rainbow Blvd., MS #3051, Kansas City, KS 66160, USA,; Pamela Tran, PhD, Department of Cell Biology and Physiology, University of Kansas Medical Center, 3901 Rainbow Blvd., MS #3038, Kansas City, KS 66160, USA. Department of Orthopaedic Surgery, University of Connecticut Health Center, Farmington, CT.

## Abstract

Cilium formation and actin cytoskeleton dynamics are interconnected, with evidence showing that elevated filamentous actin (F-actin) negatively regulates primary cilia length. Loss of the cytoskeletal protein SPECC1L, which itself does not localize to cilia, leads to increased F-actin and shortened cilia. Depolymerizing F-actin in *Specc1l* mutant cells restored cilia lengths, substantiating this inverse relationship. In cells harboring a *Specc1l* allele lacking only the coiled-coil domain 2, intracellular regions with both elevated and reduced F-actin were observed together with cilia shortening. Notably, F-actin was decreased at the ciliary base, suggesting that a different F-actin subpopulation contributes to the inverse relationship. We also identified a genetic interaction between *Specc1l* and *Thm1*, which encodes an intraflagellar transport-A (IFT-A) protein. Double or compound heterozygotes for *Specc1l* and *Thm1* exhibited a higher penetrance of cleft palate compared to *Specc1l* heterozygotes alone. Together, these findings reveal a role for SPECC1L in cytoskeletal regulation of ciliogenesis affecting palate development.

## Introduction

Cilia are microtubule-based projections that extend from the surface of nearly all vertebrate cells. These antenna-like structures are assembled and maintained through intraflagellar transport (IFT), which is the bi-directional movement of structural and signaling proteins along the microtubular axoneme (Kozminski et al. 1993). Defects in ciliary structure and function give rise to ciliopathies, which are pleiotropic disorders often associated with neural tube defects, such as exencephaly, and craniofacial abnormalities, including cleft palate (Andreu-Cervera, Catala, and Schneider-Maunoury 2021; Rosenbaum and Witman 2002; Waters and Beales 2011). IFT proteins assemble into two multiprotein complexes, IFT-B and IFT-A. The IFT-B complex mediates anterograde transport from the ciliary base to the tip, whereas the IFT-A complex is required for retrograde transport from the tip back to the base (Rosenbaum and Witman 2002). Studies indicate that the actin cytoskeleton can modulate ciliary structure and function (Avasthi et al. 2014; Hufft-Martinez et al. 2024; Jack et al. 2019; Kim et al. 2015; Woodford and Blanks 1989; Yeyati et al. 2017), and that actinopathies often share phenotypic overlap with ciliopathies (Hufft-Martinez et al. 2024). In general, levels of filamentous actin (F-actin) exhibit an inverse relationship with cilia formation and length (Avasthi et al. 2014; Hufft-Martinez et al. 2024; Jack et al. 2019; Kim et al. 2015; Woodford and Blanks 1989; Yeyati et al. 2017).

Mutations in *SPECC1L* (sperm antigen with calponin homology and coiled-coil domains 1-like) have been identified in both syndromic and isolated cases of clefts of the lip and/or palate (CL/P). *SPECC1L* encodes a cytoskeletal scaffolding protein that interacts with F-actin, microtubules, non-muscle myosin II, and junctional proteins (Saadi et al. 2023). These interactions are primarily mediated through two conserved domains: the coiled-coil domain 2 (CCD2), which binds microtubules, and the calponin homology domain (CHD), which binds F-actin (Goering, Wenger, et al. 2021; Hall et al. 2020; Mehta et al. 2023; Saadi et al. 2011; Wilson et al. 2016). We along with others have shown that patients with autosomal dominant *SPECC1L* variants are associated with a spectrum of structural birth defects, including hypertelorism, omphalocele and cleft palate (Bhoj et al. 2019; Kruszka et al. 2015; Saadi et al. 2023). Notably, most pathogenic variants in *SPECC1L* cluster within CCD2 or CHD (Goering, Wenger, et al. 2021; Hall et al. 2020; Saadi et al. 2023).

To model the human phenotypes, we have generated several *Specc1l* mouse mutant alleles, including a null allele lacking all SPECC1L protein expression (Δ*Ex4*) and an allele specifically lacking CCD2 (Δ*CCD2*) (Goering, Isai, et al. 2021). On a C57BL/6J background, mice homozygous for the *Specc1l*^Δ*Ex4*^ null allele did not show cleft palate (CP), but did show perinatal lethality, fused digits, and delayed palate elevation (Goering, Isai, et al. 2021). Here, we report findings for the null allele on the FVB/NJ background, where homozygous Δ*Ex4* mutants showed some penetrance of CP. In contrast to the null allele, Δ*CCD2* mutant embryos exhibited highly penetrant exencephaly, CP, ventral body wall closure defect and coloboma, consistent with a gain-of-function mechanism (Goering, Isai, et al. 2021).

At the cellular level, loss of SPECC1L resulted in a marked increase in F-actin in both tissues and cultured cells, indicating a role in F-actin turnover (Saadi et al. 2011; Wilson et al. 2016; Hall et al. 2020; Goering, Wenger, et al. 2021). In Δ*CCD2* cells, however, the actin cytoskeleton was even more disorganized than in the null mutant, consistent with a gain-of-function mechanism. This disorganization was characterized by subcellular regions with increased peripheral F-actin and decreased perinuclear F-actin (Goering, Wenger, et al. 2021). These mutants thus provide a valuable model to investigate which F-actin populations influence ciliogenesis. Here, we show that increased F-actin due to SPECC1L deficiency leads to shortened cilia, with a bulbous tip, which is typically observed in IFT-A mutants (Tran et al. 2008). Furthermore, *Specc1l* deficiency genetically interacts with deficiency of IFT-A gene, *Thm1* (*Ttc21b*). These findings reveal a previously unrecognized role for SPECC1L-mediated actin cytoskeletal regulation in ciliogenesis, and a genetic interaction with *Thm1* that converges on retrograde ciliary transport affecting palatogenesis.

## Materials and Methods

### Mouse embryo processing and histological analysis

Timed matings were performed overnight, and the presence of a vaginal plug was checked the following morning. Noon on the day a plug was detected was designated as embryonic day 0.5 (E0.5). Pregnant dams were sacrificed at the required embryonic stage in accordance with protocols approved by the Institutional Animal Care and Use Committee (IACUC). Embryos were collected, rinsed in 1× phosphate-buffered saline (PBS), and imaged using a Nikon SMZ1500 stereomicroscope. E15.5-18.5 embryos were also scored for presence or absence of cleft palate. Yolk sacs were retained for genotyping purposes, and sex determination was performed using PCR as described previously (Goering, Wenger, et al. 2021). Embryo heads were fixed overnight in 4% paraformaldehyde (PFA). For cryosection preparation, fixed heads were incubated at 4°C in 15% sucrose overnight, followed by incubation in 30% sucrose overnight until fully submerged, and then embedded in optimal cutting temperature (OCT) compound. Tissues were coronally sectioned at a thickness of 10 μm and stored at −80°C until immunofluorescence analysis.

### Mouse embryonic fibroblast isolation and culture

Mouse embryonic fibroblasts (MEFs) were isolated from E13.5 embryos. Embryos were decapitated, and limbs, organs, and internal structures were removed. The remaining tissue was finely minced using a sterile razor blade. Tissue fragments were incubated in 0.25% trypsin (Thermo Fisher, 25200056) at 37°C for 5 minutes. After trypsinization, cells were resuspended in high-glucose DMEM supplemented with 10% fetal bovine serum and 100 units/mL of penicillin/streptomycin (Corning, 30-002-CI). MEFs were plated in six-well plates and maintained at 37°C with 5% CO₂. Culture medium was replaced daily until the cells reached confluency, at which point they were cryopreserved. For immunostaining, frozen cells were thawed and seeded at the appropriate density.

### Immunofluorescence

For MEF immunostaining, cells were cultured on poly-lysine-coated glass coverslips and fixed with 2% paraformaldehyde for 10 minutes at room temperature (RT). Following fixation, cells were permeabilized with 0.1% Triton X-100 in 1× PBS for 10 minutes, washed three times with PBS, and then blocked in 10% normal goat serum (NGS; Thermo Fisher Scientific, 50062Z) for 1 hour at RT. Primary antibodies were applied and incubated overnight at 4°C. The following day, cells were washed with 1× PBS, incubated with secondary antibodies for 2 hours at RT, and subsequently stained with DAPI and/or phalloidin (F-actin stain; Cytoskeleton Inc.) for 30 minutes. Stained coverslips were mounted on slides using ProLong Gold Antifade Mountant (Thermo Fisher Scientific, P10144).

For E13.5-15.5 palate tissue sections, an antigen retrieval protocol was followed by heating slides in sodium citrate buffer (10 mM sodium citrate, 0.05% Tween 20, pH 6.0) at 96^◦^C for 10 min. Slides were washed in H_2_O and PBS, permeabilized using 0.5% Triton X-100 in 1× PBS for 30 min, washed in PBS again, then blocked in 10% NGS. Primary and secondary antibodies were incubated following the same method as described above. Images were acquired using one of the following: EVOS M7000 inverted imaging system, Leica SP8 STED 3× white light laser confocal microscope, Nikon Eclipse Ti-E with A1R confocal microscope, or Nikon CSU-W1 Spinning-disk confocal with SoRa microscope. Primary antibodies: SPECC1L N-terminus (1:500 cells, 1:250 tissue; Proteintech, 25390-1-AP), acetylated α-tubulin (1:4000; Sigma, T7451), ARL13B (1:300; Proteintech, 17711-1-AP), Acti-stain 555 phalloidin (1:140; Cytoskeleton, PHDH1-A), Actin-stain 670 phalloidin (1:140; Cytoskeleton, PHDN1-A), DAPI (5 μM), Secondary antibodies and stains: Goat anti-Rabbit Immunoglobulin G (IgG) (H + L) Alexa 488 and 594 (1:1000 cells, 1:500 tissue; Invitrogen, A-11008, A-11012), Goat anti-mouse IgG1 Alexa 488 (1:500; Invitrogen, A-21121), Goat anti-Mouse IgG1 Alexa 680 (1:500; Jackson ImmunoResearch, 115-625-205).

### Cilia length measurement

Cilia measurements were performed using images acquired primarily on a Nikon Eclipse Ti-E microscope equipped with an A1R confocal system. Z-stacks were captured with slices taken every 0.2 μm ensuring the full depth of each tissue section was captured. Maximum intensity projections (MIPs) were created for measurement. Cilia lengths were measured using ImageJ software and a segmented line tool was used to trace the cilium, following the method described by Jack & Avasthi (Jack et al. 2019). Measurements in pixels were converted to microns using the appropriate pixel-to-micron conversion factor for the objective used. Cilia length data were analyzed and graphed using GraphPad Prism, with 95% confidence intervals represented.

### Ciliogesnesis and Chemical Modulation

MEFs were cultured to approximately 70% confluence and then serum-starved by replacing the growth medium with DMEM alone for 18 hours to induce cilia formation. Following serum starvation, cells were treated for 2 hours with either DMSO (vehicle control), 0.2 mM or 2 mM Latrunculin B (Sigma, 428020). For Jasplakinolide treatment, MEFs were exposed to 200nM Jasplakinolide (Cayman Chemical, 102396-24-7) for 2 hours occurred following 18-hour serum starvation period.

### Pericentriolar Region Analysis

MEFs were cultured on #1.5 glass coverslips at high density and serum starved for 18 hours to promote primary cilia growth. Cells were then fixed in 2% PFA for 10 minutes, permeabilized using 0.1% Triton X-100 for 30 minutes and blocked overnight using 10% NGS. Cells were immunostained using anti-acetylated-α-tubulin, anti-SPECC1L, and phalloidin. High-resolution z-stack images were obtained using the Nikon CSU-W1 spinning-disk confocal with SoRa and then deconvolved using Nikon NIS-Elements software. For pericentriolar analysis, a 1 µm^3^ volume was defined at the base of the cilium so that it included the cytoplasmic region immediately beneath the ciliary axoneme. Nikon NIS-Elements analysis software was used to obtain the summed intensity of phalloidin and SPECC1L fluorescence within this 1 µm^3^ volume.

### Cell Fluorescence Quantification

Cell fluorescence intensity was quantified using ImageJ/Fiji software. The corrected total cell fluorescence (CTCF) was calculated using this formula (Ansari et al. 2013): CTCF=Integrated Density -(Area of selected area X Mean fluorescence of background readings).

For measurements, “Area”, “Integrated Density”, and “ Mean Grey value” were selected under *Set Measurements*. Three background regions were measured to obtain an average background intensity. The region on interest (ROI) was then measured to obtain the integrated density value, and CTCF values were calculated accordingly. Final values were graphed using GraphPad Prism, with data represented as the mean ± standard deviation (SD).

## Results

### Specc1l mutants have shortened primary cilia in vitro and in vivo

SPECC1L is not present in primary cilia, but colocalizes with F-actin in the cytoplasm. Further, the loss of SPECC1L leads to an increase in F-actin (Saadi et al. 2011; Wilson et al. 2016; Hall et al. 2020; Goering, Wenger, et al. 2021). Since F-actin has been shown to regulate ciliogenesis (Avasthi et al. 2014; Hufft-Martinez et al. 2024; Jack et al. 2019; Kim et al. 2015; Lee et al. 2018; Woodford and Blanks 1989; Yeyati et al. 2017), we hypothesized that *Specc1l* deficiency may impair ciliogenesis. To examine cilia, we immunostained E13.5 mouse craniofacial tissue sections with acetylated-α-tubulin (axoneme marker) and ARL13B (ciliary membrane marker) (**Figure 1A-C**). In E13.5 palate mesenchyme from both *Specc1l*^Δ*Ex4/*Δ*Ex4*^ null and *Specc1l*^Δ*CCD2/*Δ*CCD2*^ gain-of-function mutants lacking CCD2, cilia lengths were reduced (**Figure 1A-C**). The reduction was similar in null Δ*EX4* and Δ*CCD2* mutants (**Figure 1D**), indicating that disruption of SPECC1L-microtubule association and the resulting peripheral F-actin accumulation was sufficient to impair ciliogenesis.

**Figure 1.**
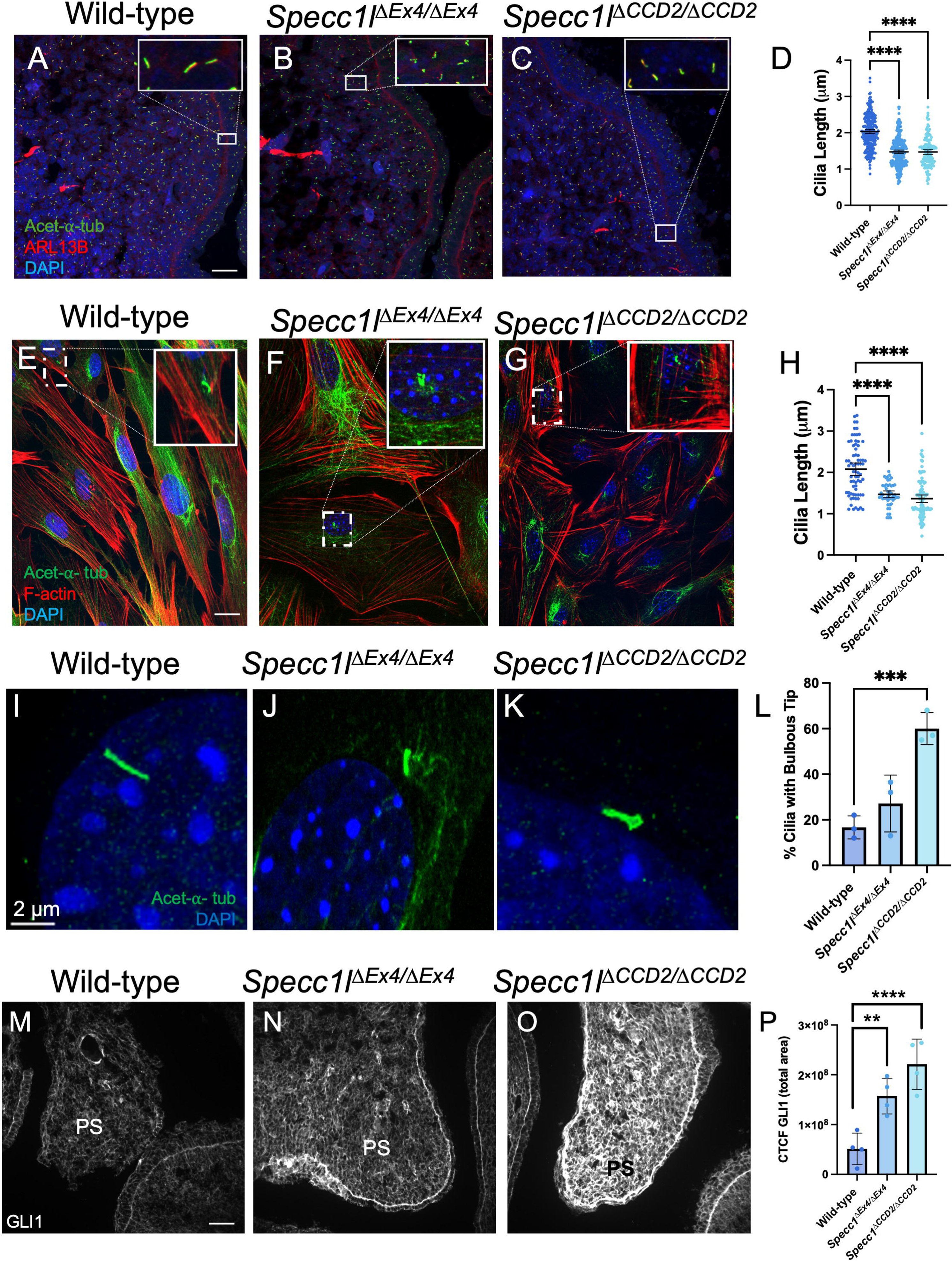
*Specc1l* mutants have shortened primary cilia *in vitro* and *in vivo*. **A-C.** Embryonic day (E) 13.5 mouse palate mesenchyme tissue from wildtype (WT) **(A),** *Specc1l*^Δ*Ex4/*Δ*Ex4*^ null **(B),** and *Specc1l*^Δ*CCD2/*Δ*CCD2*^ **(C)** embryos, stained with acetylated-α-tubulin (green), ARL13B (red), and DAPI. The insets show 2x zoom of select regions. **D.** Quantification of cilia length from 3 different embryos of each genotype (WT, n=250; ΔEx4, n=290; ΔCCD2, n=200, **** <0.0001, Scale bar represents 2μm**). E-G.** Mouse embryonic fibroblasts (MEFs) from WT **(E)**, *Specc1l*^Δ*Ex4/*Δ*Ex4*^ **(F)**, and *Specc1l*^Δ*CCD2/*Δ*CCD2*^ **(G)** embryos stained with acetylated-α-tubulin (green), F-actin (red), and DAPI. **H.** Quantification of cilia length from 3 different experiments (n=150 cilia/genotype, **** <0.0001, Scale bar represents 2μm**). I-L.** MEFs from WT **(I),** *Specc1l*^Δ*Ex4/*Δ*Ex4*^ **(J),** and *Specc1l*^Δ*CCD2/*Δ*CCD2*^ **(K)** stained with acetylated-α-tubulin (green) and imaged at 100x with 4x zoom revealed bulbous tips in *Specc1l*^Δ*CCD2/*Δ*CCD2*^ **(K). L.** Quantification of bulbous tips from 3 different experiments (n=150 cilia/genotype, *** <0.001, Scale bar represents 2μm). **M-O.** E13.5 mouse palate mesenchyme tissue from WT **(M),** *Specc1l*^Δ*Ex4/*Δ*Ex4*^ **(N),** and *Specc1l*^Δ*CCD2/*Δ*CCD2*^ **(O)** stained with GLI1 (white). **P.** Quantification of GLI1 from 3 different embryos of each genotype (** <0.0044, **** <0.0001, Scale bar represents 2μm, Error bars represent 95% CI for (D &H) and Mean ± standard deviation (SD) for (L &P)).

For a closer examination, mouse embryonic fibroblasts (MEFs) were harvested from Δ*EX4* and Δ*CCD2* mutant E13.5 embryos. Cilia lengths were reduced in MEFs from both *Specc1l* mutants compared to wild-type (**Figure 1E-H**). Additionally, super-resolution microscopy revealed *Specc1l*^Δ*CCD2/*Δ*CCD2*^ mutant cells have a statistically significant increase in cilia with bulbous distal tips (**Figure 1I-L**), a hallmark of retrograde IFT-A mutants like *Thm1* (Duran et al. 2017; Fu et al. 2016; Jacobs et al. 2016; Jacobs et al. 2020; Miller et al. 2013; Qin et al. 2011; Tran et al. 2008; Tran et al. 2014; Wang et al. 2020).

We next assessed whether the ciliary shortening had functional consequences by examining levels of GLI1, a downstream effector of cilia-dependent Hedgehog signaling (Yin et al. 2022; Tran et al. 2008; Tran et al. 2014). GLI1 expression was significantly increased in E13.5 palate tissues from both *Specc1l* mutant alleles (**Figure 1M-P**), indicating altered ciliary function.

### Depolymerization of F-actin restores length of Specc1l mutant shortened primary cilia

To test whether cilia shortening in *Specc1l* mutant cells resulted from increased actin filament stability, we treated MEFs with 200nM jaskplakinolide, which stabilizes actin filaments by inducing nucleation (Bubb et al. 2000). Indeed, jaskplakinolide treatment shortened cilia in wild-type cells but had no additional effect on *Specc1l* mutant cells (**Figure 2A**), suggesting that actin filaments in mutant cells were already extensively stabilized, consistent with an inverse relationship between actin stability and cilia length.

**Figure 2.**
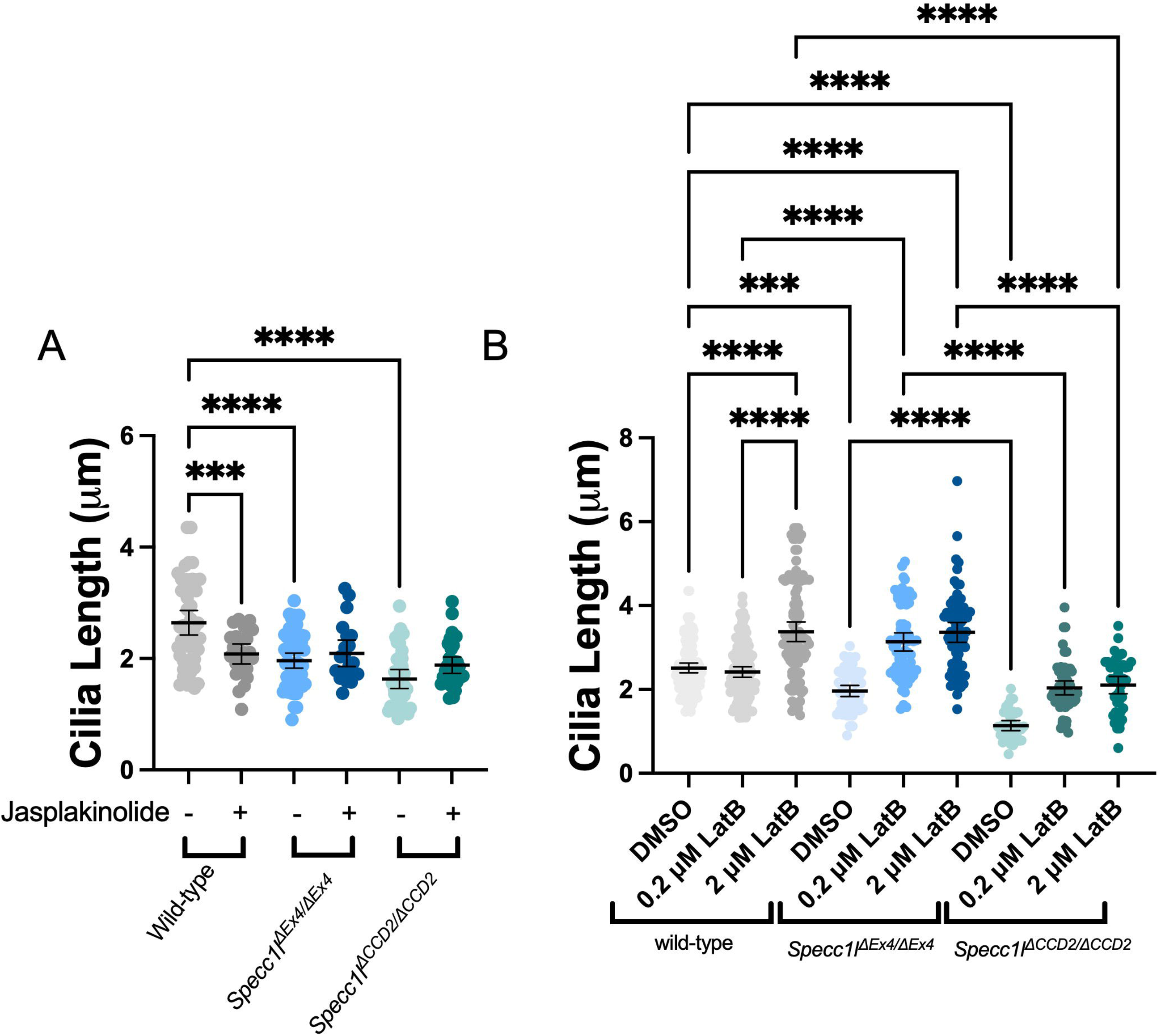
Chemical depolymerization of F-actin rescues cilia length in *Specc1l* mutants. **A.** Quantification of primary cilia length in mouse embryonic fibroblasts (MEFs) from WT, *Specc1l*^Δ*Ex4/*Δ*Ex4*^, and *Specc1l*^Δ*CCD2/*Δ*CCD2*^ treated with DMSO (vehicle) and Jasplakinolide (n= 65/genotype, *** <0.0004, ****<0.0001). **B.** Quantification of primary cilia length in WT, *Specc1l*^Δ*Ex4/*Δ*Ex4*^, and *Specc1l*^Δ*CCD2/*Δ*CCD2*^ MEFs treated with DMSO and Latrunculin B (LatB) (n= 70/genotype, *** 0.0006, ****<0.0001, Error bars represent 95% CI). Data are shown from 3 different experiments.

To further test the relationship between actin dynamics and cilia length, we asked whether F-actin depolymerization could restore normal cilia length in mutant cells. MEFs were treated with Latrunculin B (LatB), which inhibits actin polymerization by binding to G-actin monomers, at 0.2 μM or 2 μM LatB for 2 hours (Jack et al. 2019). As expected, in wild-type MEFs treated with the higher 2 μM concentration, cilia increased in length from an average of 2.5 microns (μm) to 3 μm. In *Specc1l*^Δ*Ex4/*Δ*Ex4*^ null mutant MEFs, cilia lengths reached ∼3 μm length with just 0.2 μM LatB treatment and maintained this length at 2 μM LatB. In contrast, *Specc1l*^Δ*CCD2/*Δ*CCD2*^ mutant cilia reached only ∼2.5 μm length at 0.2 μM LatB, with no further increase at 2 μM LatB (**Figure 2B, Figure S1**). These results indicate that *Specc1l* mutant cells are more sensitive to actin depolymerization, and that the difference in cilia length rescue between null *ΔEx4* and gain-of-function *ΔCCD2* mutant cells may reflect the severity of actin cytoskeletal defects, with *ΔCCD2* mutant cells being more severely affected (Goering, Wenger, et al. 2021).

### Filamentous actin is reduced in the pericentriolar region of SPECC1L mutant cells

We previously showed that in addition to F-actin, WT-SPECC1L localized to the pericentriolar region near gamma-tubulin, a centriolar marker (Saadi et al. 2011). Using super-resolution microscopy, we examined SPECC1L and F-actin localization at the base of the cilium in both WT and *Specc1l*^Δ*CCD2/*Δ*CCD2*^ mutant MEFs (**Figure 3A, B**). Analysis of a 1 μm^3^ cubic region around the cilium base revealed reduced SPECC1L-ΔCCD2 mutant protein and decreased F-actin in mutants (**Figure 3C, D**). This reduction of F-actin at the ciliary base in *Specc1l* mutant cells suggested that the increased F-actin driving cilia shortening originated from an actin population elsewhere in the cell.

**Figure 3.**
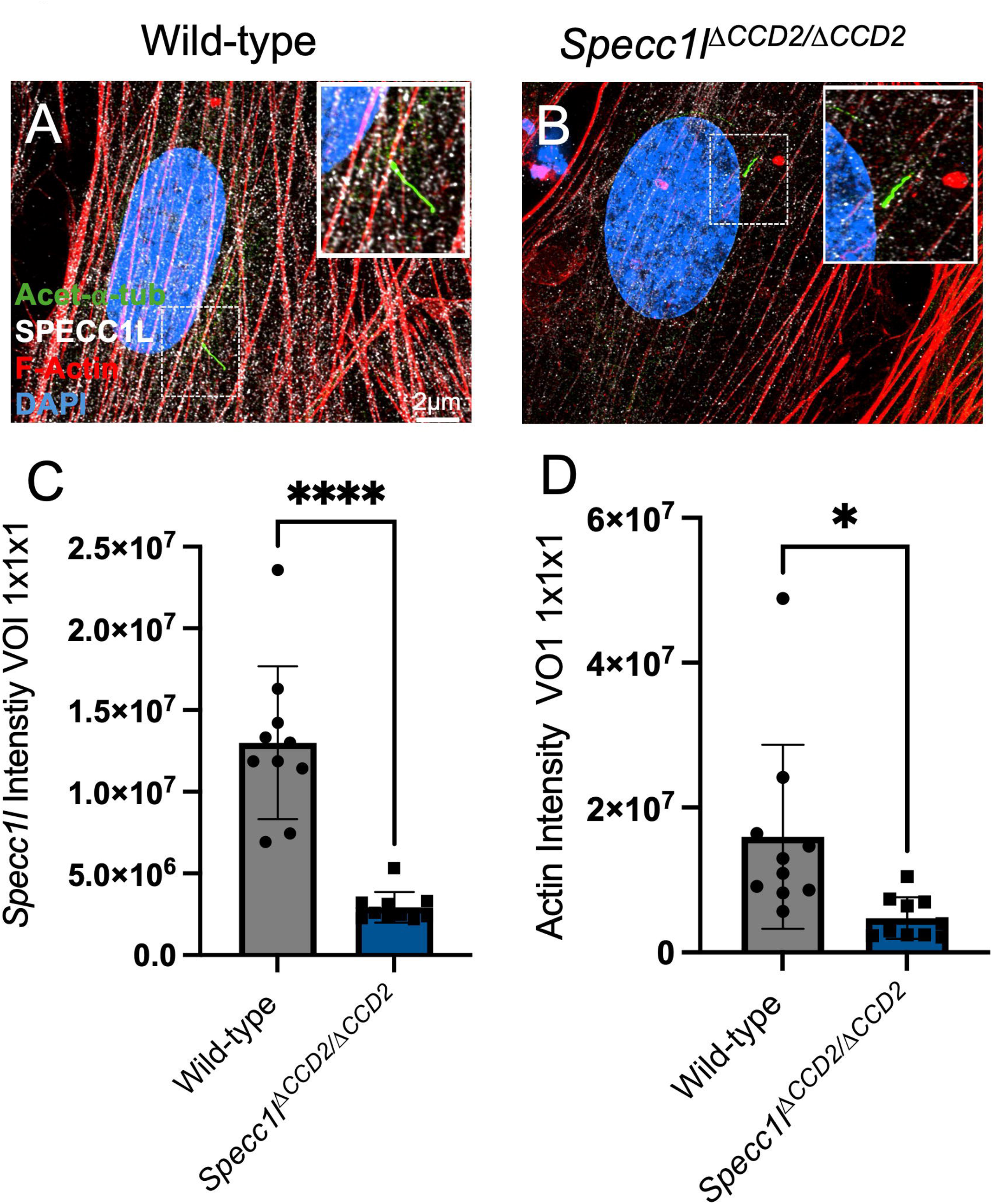
*Specc1l*^Δ*CCD2/*Δ*CCD2*^ display decreased SPECC1L and F-actin expression in the pericentriolar region. **A-B.** Mouse embryonic fibroblasts (MEFs) from WT**(A)** and *Specc1l*^Δ*CCD2/*Δ*CCD2*^ **(B)** were stained for SPECC1L (white), F-actin (red), acetylated-α-tubulin (green) and DAPI. **C-D.** Quantification of a 1×1x1 µm cubic volume of interest (VOI) for SPECC1L **(C)** and F-actin **(D)** showed a significant reduction for both in ΔCCD2 mutant cells (****<0.0001, * 0.0138, N=3, n=15 cilia, Scale bar represents 2μm, Error bars represent Mean ± standard deviation (SD)).

### Cytoskeletal Specc1l and ciliary Thm1 genetically interact

Since shortened cilia with bulbous distal tips observed in *Specc1l* mutant cells (**Figure 1**) resembles an IFT-A mutant ciliary phenotype, we tested for a genetic interaction with IFT-A gene, *Thm1* (or *Ttc21b*). We used the *Thm1* aln allele which carries a missense mutation and results in loss of protein, effectively making it a null allele. Homozygous *Thm1^aln/aln^* mutants exhibit exencephaly, cleft palate, and shortened bulbous cilia (Tran et al. 2008). We crossed *Specc1l* (*null* Δ*Ex4 or* Δ*CCD2*) and *Thm1* (*aln*) single heterozygous mice on an FVB/NJ background. *Specc1l*^Δ*Ex4/+*^ heterozygotes did not show cleft palate (**Figure 4A**, n=46), consistent with previous observation on a C57BL/6J background (Goering, Wenger, et al. 2021). *Specc1l*^Δ*Ex4/+*^*;Thm^aln/+^*double heterozygous embryos showed a single occurrence of CP (**Figure 4A**, n=28). *Specc1l*^Δ*Ex4/*Δ*Ex4*^ homozygous mutants displayed ∼21% CP (**Figure 4A-E**, n=37) on the FVBN/J background, validating SPECC1L’s role in palatogenesis (Goering, Wenger, et al. 2021). Notably, *Specc1l*^Δ*Ex4/*Δ*Ex4*^*;Thm^aln/+^*compound mutants exhibited a significantly higher ∼58% penetrance of CP (n=12) and more severe digit fusion defects (**Figure 4A-E**), demonstrating a genetic interaction between *Specc1l* and *Thm1*.

**Figure 4.**
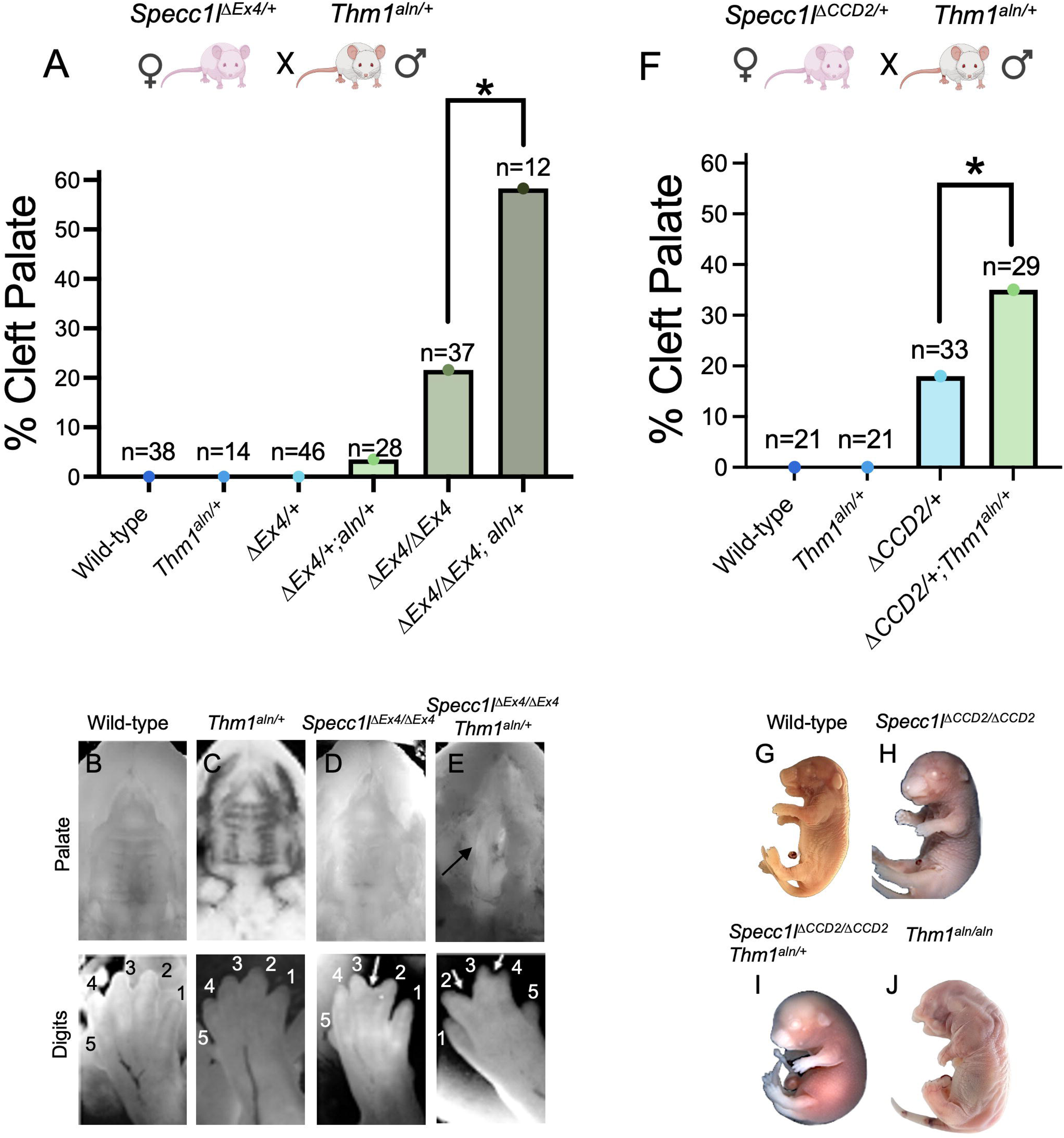
Cytoskeletal SPECC1L and ciliary THM1 genetically interact and affect palatogenesis. **A.** *Specc1l*^Δ*Ex4/+*^ single heterozygotes did not exhibit cleft palate (CP), while 10% of *Specc1l*^Δ*Ex4/*Δ*Ex4*^ homozygous mutants showed CP. Double heterozygous *Specc1l*^Δ*Ex4/+*^*; Thm1^aln/+^* embryos displayed 21.6% CP penetrance, and *Specc1l*^Δ*Ex4/*Δ*Ex4*^*; Thm1^aln/+^* compound mutants exhibited 58.3% CP, consistent with a genetic interaction (** <0.002, z-statistic, Error bar represents Mean ± standard deviation (SD)). **B-E**. E17.5 palate (top) and forelimb digit (bottom) images from WT (**B**), *Thm1^aln/+^* (**C**), *Specc1l*^Δ*Ex4/*Δ*Ex4*^ (**D**), *Specc1l*^Δ*Ex4/*Δ*Ex4*^*; Thm1^aln/+ +^*(**E**), showing worsened digit fusion in compound mutants (**D vs. E**). **F.** *Specc1l*^Δ*CCD2/+*^ single heterozygotes showed 18% CP, which increased to 35% CP in *Specc1l*^Δ*CCD2/+*^*;Thm1^aln/+^* double heterozygous embryos, consistent with a genetic interaction (* <0.017, z-statistic, Mean ± standard deviation (SD)). **G-J**. WT (**G**) and *Specc1l*^Δ*CCD2/*Δ*CCD2*^ (**H**) embryos did not show any gross abnormalities at E18.5. In contrast, the *Specc1l*^Δ*CCD2/*Δ*CCD2*^;*Thm1^aln/+^*compound mutant (**I**) showed a worsened phenotype with edema and craniofacial anomalies similar to those observed in *Thm1^aln/aln^* mutant embryo (**J**).

For the *Specc1l*^Δ*CCD2*^ allele, ∼18% of heterozygous embryos on the FVB/NJ background showed CP (**Figure 4F**, n=33), higher than the ∼10% previously reported on the C57BL/6J background (Goering, Wenger, et al. 2021). Strikingly, ∼35% of *Specc1l*^Δ*CCD2/+*^*;Thm^aln/+^*double heterozygous embryos displayed CP (**Figure 5F**, n=29), indicating a strong genetic interaction. *Specc1l*^Δ*CCD2/*Δ*CCD2*^ homozygous mutant embryos at E18.5 exhibited ∼60% cleft palate and ∼40% exencephaly, and did not show other defects (**Figure 4G, H**) (Goering, Wenger, et al. 2021). In contrast, *Specc1l*^Δ*CCD2/*Δ*CCD2*^*;Thm^aln/+^*compound mutants were more severely affected, showing extensive edema and severe craniofacial abnormalities (**Figure 4H, I**) that were more similar to craniofacial defects of *Thm1^aln/aln^* homozygous mutant embryos (**Figure 4J**), further supporting a genetic interaction.

**Figure 5.**
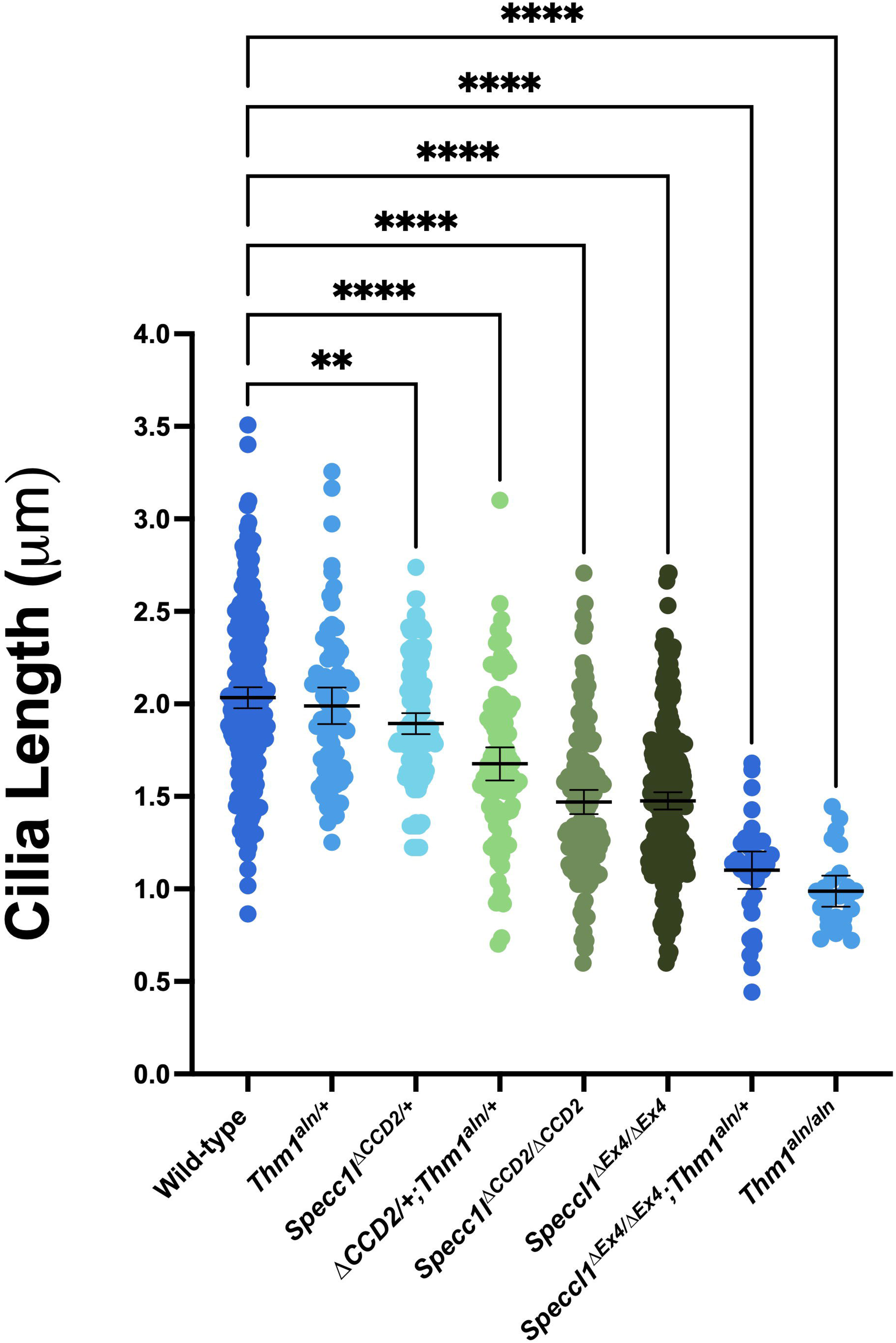
Primary cilia length shortening correlates with phenotypic severity. Quantification of primary cilia length was performed on palate mesenchyme tissue from three (E) 13.5 embryos with WT (n=250), *Thm1^aln/+^* (n=75)*, Specc1l*^Δ*CCD2/+*^ (n=150)*, Specc1l*^Δ*CCD2/+*^*;Thm1^−/+^* (n=90)*, Specc1l*^Δ*CCD2/*Δ*CCD2*^ (n=150)*, Specc1l*^Δ*Ex4/*Δ*Ex4*^ (n=290), *Specc1l*^Δ*Ex4/*Δ*Ex4*^*;Thm1^aln/+^* (n=50), and *Thm1^aln/aln^* (n=50) genotypes (** <0.0078, **** <0.0001, Error bars represent 95% CI).

Consistent with the genetic interaction, cilia length decreased in an allelic dose-dependent manner, with the mildest reduction observed in *Specc1l*^Δ*CCD2/+*^ heterozygotes, moderate reductions in *Specc1l*^Δ*CCD2*^*;Thm^aln/+^*double heterozygotes and *Specc1l* homozygous mutants (*Specc1l*^Δ*Ex4/*Δ*Ex4*^ and *Specc1l*^Δ*CCD2/*Δ*CCD2*^), and severe reduction in *Specc1l*^Δ*Ex4/*Δ*Ex4*^*;Thm^aln/+^*compound mutants. The *Thm1^aln/aln^* homozygous mutant cilia were observed to be the shortest (**Figure 5**).

### Actin Cytoskeleton defects in Thm1 mutant cells

In *Specc1l* mutant cells, the primary defect was increased actin stability inversely affecting cilia length. We asked whether the ciliary defect in *Thm1* mutants influenced the actin cytoskeleton. *Thm1^aln/aln^* null MEFs showed reduced F-actin staining (**Figure 6 A-C**). To confirm this, we quantified cell circularity (width/length), since decreased F-actin should result in a more circular cell. Consistently, *Thm1^aln/aln^* null cells were more circular than controls (**Figure 6D**).

**Figure 6.**
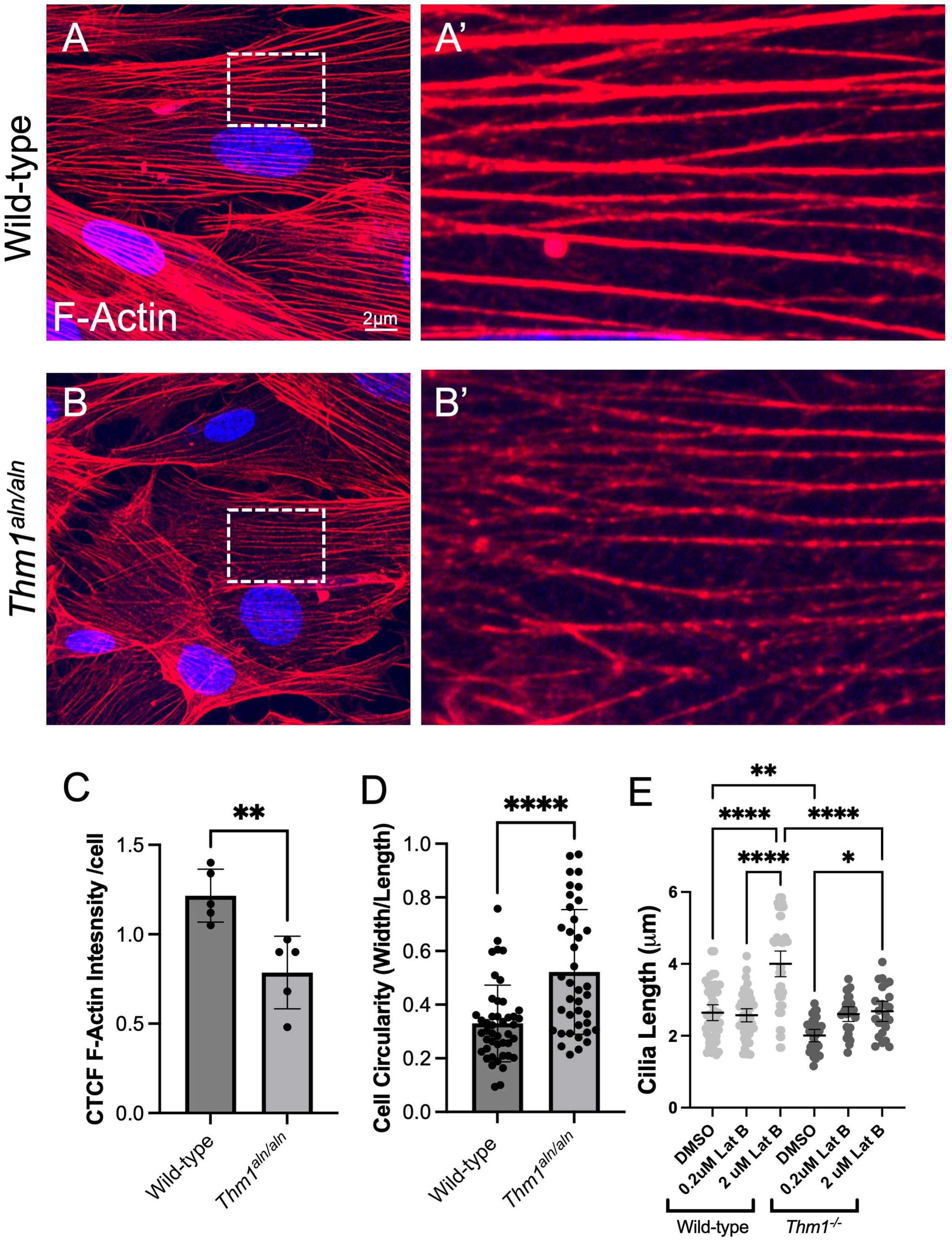
Cytoskeletal feedback in *Thm1* mutant cells. **A-B**. Mouse embryonic fibroblasts (MEFs) from WT**(A)** and *Thm1^aln/aln^***(B)** embryos were stained with F-actin (red) and DAPI (blue). **A’-B’.** 2x zoom of the boxed regions of cells shown in **A-B**. **C**. Quantification of mean gray value for actin/cell from 5 experiments (** <0.0051). **D**. Quantification of cell circularity ratio (width/length) from 3 experiments (n=45/genotype, ****<0.0001). **E**. Quantification of primary cilia length in WT and *Thm1^aln/aln^* MEFs treated with vehicle (DMSO) and Latrunculin B (LatB) from 3 experiments (n= 50, *<0.0234, **<0.0071, ****<0.0001, Scale bar represents 2μm, Error bars represent 95% CI).

Next, we investigated whether F-actin depolymerization with LatB could rescue ciliary length in *Thm1* mutants. *Thm1^aln/aln^* cells also responded to the lower 0.2 μM dose of LatB, increasing their cilia lengths to an average length of 2.5 μm from 2 μm (**Figure 6E**). The higher 2 μM dose did not further increase *Thm1* mutant cilia lengths, and these cells did not reach the maximal WT cilia lengths. Taken together, these results indicate that *Thm1^aln/aln^* null cells have limited capacity to respond to actin cytoskeletal modulation, likely due to already reduced levels of F-actin, suggesting feedback from shortened primary cilia to the actin cytoskeleton.

## Discussion

In this study, we identified a novel role for non-ciliary cytoskeletal scaffolding protein SPECC1L in ciliogenesis. SPECC1L promotes F-actin turnover, and its deficiency leads to increased cellular F-actin. We showed that *Specc1l* mutant cells exhibit shortened primary cilia, often with a bulbous distal tip, which is a hallmark of disrupted retrograde IFT. Our findings in an *in vivo* model confirm and extend previous reports of an inverse relationship between F-actin and cilia length (Avasthi et al. 2014; Jack et al. 2019; Hufft-Martinez et al. 2024; Kim et al. 2015; Lee et al. 2018; Woodford and Blanks 1989; Yeyati et al. 2017; Drummond 2012).

We validated this inverse relationship using pharmacologic manipulation of actin dynamics. Treatment with jasplakinolide, which stabilizes F-actin (Bubb et al. 2000), did not further shorten cilia in *Specc1l* mutant cells, suggesting that ciliary shortening had reached a plateau due to pre-existing F-actin stabilization. In contrast, depolymerization of F-actin with latrunculin B (LatB) at low concentrations rapidly restored cilia length, confirming that increased F-actin was a causal factor. Notably, this rescue was complete in *Specc1l*^Δ*Ex4*^ null mutant cilia, but only partial in *Specc1l*^Δ*CCD2*^ mutant cilia. This difference in response is consistent with our prior gain-of-function model for ΔCCD2 (Goering, Wenger, et al. 2021), where additional cytoskeletal changes beyond increased F-actin may be at play.

Another key finding stemmed from the fact that both *Specc1l*^Δ*Ex4*^ null and *Specc1l*^Δ*CCD2*^ mutant cells exhibited shortened cilia. In *Specc1l* null cells, F-actin is uniformly increased, whereas Δ*CCD2* cells show heterogenous F-actin distribution (Goering, Wenger, et al. 2021). Removal of CCD2 in-frame prevents SPECC1L-ΔCCD2 protein from associating with microtubules, causing it to accumulate around the nucleus and not reach the cell periphery. However, since SPECC1L-ΔCCD2 mutant protein can still bind and turn over F-actin, this abnormal subcellular expression results in reduced F-actin centrally and increased F-actin at the cell periphery (Goering, Wenger, et al. 2021). Given that mutants for both alleles showed LatB-responsive cilia shortening and ΔCCD2 cells showed reduced F-actin near the ciliary base, we propose that peripheral F-actin accumulation drives the cilia shortening (**Figure 7**). Together, these findings support a working model in which SPECC1L regulates actin organization to maintain proper ciliary length and function (**Figure 7A**). In this model, loss of SPECC1L-CCD2 leads to increased F-actin toward the cell periphery, likely disrupting the microtubule-actin interface near the ciliary base and impairing retrograde IFT (**Figure 7B**).

**Figure 7.**
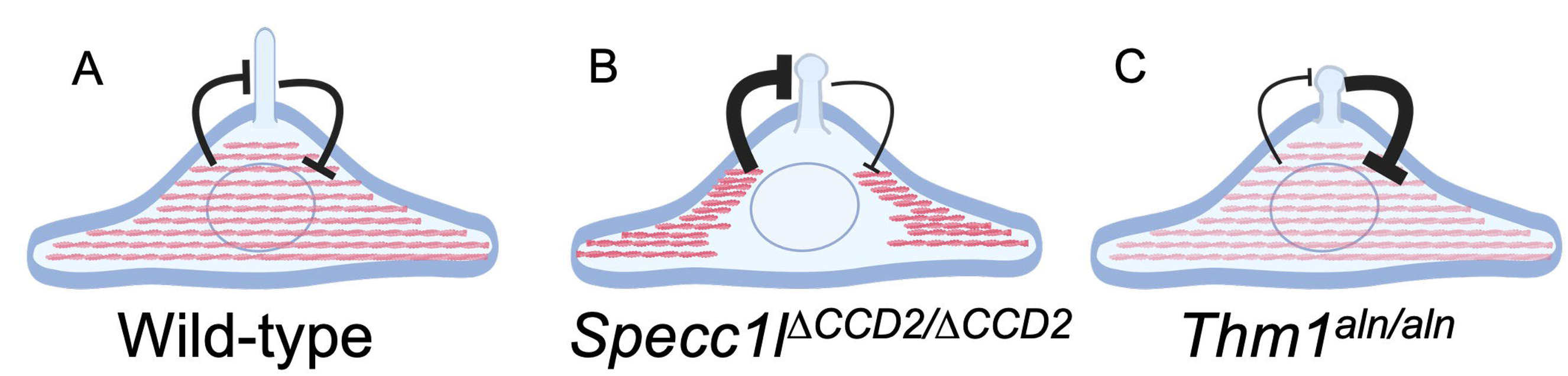
Cytoskeletal and Ciliary Feedback Model. **A-C**. Schematic representation of mesenchymal cell, highlighting the actin cytoskeleton and primary cilium. **A**. Wild-type cell with normal distribution of actin cytoskeleton and primary cilia in homeostasis. **B**. *Specc1l*^Δ*CCD2/*Δ*CCD2*^ cell with abnormal distribution of the actin cytoskeleton including a halo around the nucleus. Distribution of the actin cytoskeleton in the cytoplasm has increased inhibition on ciliogenesis. **C**. *Thm1^aln/aln^* cell showing decreased intensity of the F-actin cytoskeleton and decreased inhibition on the primary cilium, resulting from the IFT-A defect.

The genetic interaction between *Specc1l* alleles (null or Δ*CCD2*) and IFT-A mutant, *Thm1^aln^*, indicated that actin cytoskeletal disorganization and a retrograde IFT defect can converge to affect ciliary length. While *Specc1l^ΔCCD2/^*^Δ*CCD2*^ mutant embryos exhibit multiple tissue movement and fusion defects, including exencephaly, cleft palate, omphalocele and coloboma (Goering, Wenger, et al. 2021), the double heterozygotes mainly manifested CP phenotype. This result highlights the particular sensitivity of palate closure to the combined effects of actomyosin disorganization and ciliary dysfunction.

Little is known about how ciliary defects influence the actin cytoskeleton. To begin to explore this relationship, we examined F-actin in IFT-A (*Thm1*) deficient MEFs and observed reduced F-actin staining and a more rounded cell morphology. Furthermore, LatB treatment could only partially rescue *Thm1* null cilia length. These findings may suggest a feedback loop in which shortened cilia signal back to the cytoskeleton, reducing F-actin to maximize cilia length (**Figure 7C**).

Our studies establish a mechanistic connection between actomyosin regulation, ciliary architecture, and tissue morphogenesis, providing new insights into how cytoskeletal perturbations contribute to ciliopathies and craniofacial developmental disorders.

## Resource availability

### Lead contact

Requests for further information and resources and reagents should be directed to the lead contact, Irfan Saadi.

### Materials availability

This study did not generate new unique reagents.

### Data and code availability

- All data are available in the main text.
- This paper does not report any original code.
- Any additional information required to reanalyze the data reported in this paper is available from the lead contact upon request.

## Acknowledgments

This project was supported in part by the National Institutes of Health grants DE026172, DE032825 (IS), DE032515, DE032742 (IS, PT), and F31DE031181, TL1R002368 (BMH). The project was also supported in part by the Center of Biomedical Research Excellence (COBRE) grant (National Institute of General Medical Sciences P30 GM122731), Kansas IDeA Network for Biomedical Research Excellence grant (National Institute of General Medical Sciences P20 GM103418), and Kansas Intellectual and Developmental Disabilities Research Center (KIDDRC) grant (Eunice Kennedy Shriver National Institute of Child Health and Human Development, U54 HD090216). The Leica STED microscope was supported by NIH S10 OD023625. The Nikon CSU-W1 SoRa microscope was supported by NIH S10 OD032207.

## Author contributions

B.M.H, J.P.G., P.V.T. and I.S. conceived and designed the experiments. B.M.H., J.P.G., M.S., M.M., A.T., D.N.T., S.C.W., and H.H.W. performed the experiments. B.M.H., P.V.T., and I.S. wrote the paper. D.N.T., A.T., J.P.G., and M.S. edited the manuscript. All authors reviewed the manuscript.

## Supplemental information

**Figure S1. Treatment with LatB rescues cilia length in mutant MEFs**

