## Supplemental Figure 1 for "Genetic interaction of *Specc1l* and *Thm1* reveals cytoskeletal–ciliary crosstalk"

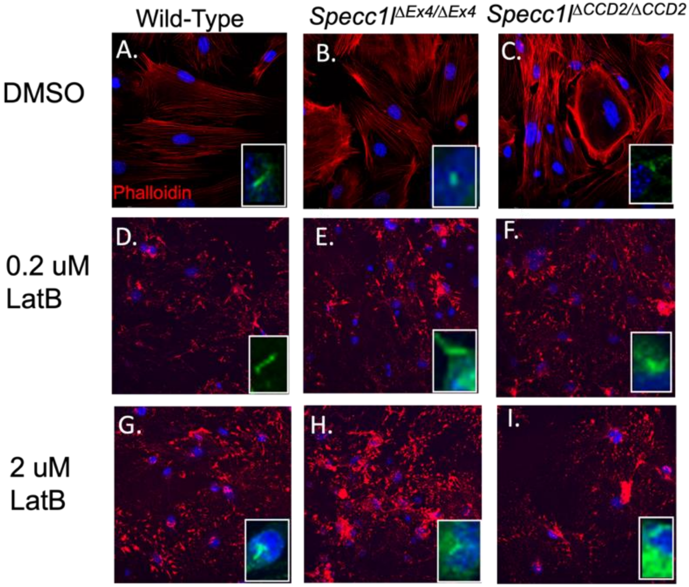


**Figure S1. Treatment with LatB rescues cilia length in mutant MEFs**

**A-C.** Mouse embryonic fibroblasts (MEFs) treated with DMSO for two hours, stained with Phalloidin (red) with cilia inset stained with acetylated-$\alpha$-tub wild-type (**A**), *Specc1l*$^{\Delta Ex4/\Delta Ex4}$ (**B**), *Specc1l*$^{\Delta CCD2/\Delta CCD2}$(**C**). **D-E.** MEFs treated with 0.2mM LatB for two hours, stained with Phalloidin (red) with cilia inset stained with acetylated-$\alpha$-tub wild-type (**D**), *Specc1l*$^{\Delta Ex4/\Delta Ex4}$ (**F**), *Specc1l*$^{\Delta CCD2/\Delta CCD2}$(**F**). **G-I.** MEFs treated with 2mM LatB for two hours, stained with Phalloidin (red) with cilia inset stained with acetylated-$\alpha$-tub wild-type (**G**), *Specc1l*$^{\Delta Ex4/\Delta Ex4}$ (**H**), *Specc1l*$^{\Delta CCD2/\Delta CCD2}$(**I**).
